# Detergent wash improves vaccinated lymph node handling ex vivo

**DOI:** 10.1101/2020.09.07.286393

**Authors:** Alexander G. Ball, Maura C. Belanger, Rebecca R. Pompano

## Abstract

Lymph nodes (LNs) are essential secondary immune organs where the adaptive immune response is generated against most infections and vaccines. We recently described the use of live ex vivo LN slices to study the dynamics of adaptive immunity. However, when working with reactive lymph nodes from vaccinated animals, the tissues frequently became dislodged from the supportive agarose matrix during slicing, leading to damage that prevented downstream analysis. Because reactive lymph nodes expand into the surrounding adipose tissue, we hypothesized that dislodging was a result of excess lipids on the collagen capsule of the LN, and that a brief wash with a mild detergent would improve LN interaction with the agarose without damaging tissue viability or function. Therefore, we tested the use of digitonin on improving slicing of vaccinated LNs. Prior to embedding, LNs were quickly dipped into a digitonin solution and washed in saline. Lipid droplets were visibly removed by this procedure. A digitonin wash step prior to slicing significantly reduced the loss of LN during slicing from 13 - 75 % to 0 - 25 %, without substantial impact on viability. Capture of fluorescent microparticles, uptake and processing of protein antigen, and cytokine secretion in response to a vaccine adjuvant, R848, were all unaffected by the detergent wash. This novel approach will enable ex vivo analysis of the generation of adaptive immune response in LNs in response to vaccinations and other immunotherapies.

## 1. Introduction

Lymph nodes (LN) are critical secondary lymphoid organs that initiate much of adaptive immunity. We recently reported an optimized approach for the collection and use of live slices of murine LN to study the development of the adaptive immune response ex vivo.^1^ In this model, skin-draining LNs are harvested, embedded in low-melting agarose, sliced on a vibratome, and then studied directly or cultured overnight. The LN slice system provides information on spatial organization and dynamics within LNs, while simultaneously allowing quantification of cytokine secretion and other in vitro assay methods.

Although this platform has the potential to enable the study of responses to vaccines and other immunotherapies in LNs undergoing an adaptive immune response,^1,2^ previous work with LN slices for T cell motility worked only with tissues from naïve animals.^3,4^ Unexpectedly, collection of slices from immunologically reactive LNs from vaccinated animals proved challenging. Specifically, within a few days of in vivo vaccination, harvested lymph nodes were frequently dislodged or “popped out” from the agarose during slicing. Lymph nodes popping out of the agarose severely reduced the number of slices that could be analyzed, as a partially sectioned LN is damaged and unusable for further slicing. Furthermore, selective loss of the most reactive LNs may prevent accurate ex vivo analysis of interesting immunological events such as germinal center formation and inflammatory processes.

To address this problem, we considered the fact that after vaccination, reactive LNs expand into the surrounding adipose tissue and potentially interact with the lipids there.^5,6^ Therefore, we hypothesized that popping could be prevented by removing lipids from the outer LN capsule, by washing in detergent before slicing. Our goal was to solubilize any lipids adsorbed to the proteinaceous, collagen-rich LN capsule, without disrupting the viability or function of the cells beneath. Therefore, we sought a mild detergent whose diffusion through the capsule would be slow.

Properties such as charge and size greatly influence a detergent’s ability to remove lipids, biocompatibility (how quickly it denatures plasma membranes and proteins), and transport properties.^7^ Non-ionic detergents are milder than ionic detergents, in that they spare protein-protein interactions while disrupting lipid-lipid and lipid-protein interactions.^8^ Detergents with a large molecular weight should be hindered from diffusing through the tightly packed LN capsule. Based on these criteria, we identified digitonin as a promising detergent that with the potential to preserve LN slice viability. Digitonin is a weak, non-ionic detergent with a complex polysaccharide headgroup and a cholesterol derivative tail, does not destroy plasma membranes, and has a relatively large molecular weight for a surfactant (1229.31 g/mol).^9,10^

Here, we tested a digitonin washing step on whole skin-draining LNs to improve the reproducibility of slicing of vaccinated LNs. In addition to the frequency of popping from the agarose, we assessed digitonin’s impact on LN slice viability in both vaccinated and naïve slices and the detergent’s impact on tissue functions such as stimulated cytokine secretion, particle capture, and antigen uptake and processing.

## 2. Materials and Methods

### 2.1 Lymph Node Slicing

All animal work was approved by the Institutional Animal Care and Use Committee at the University of Virginia under protocol #4042, and was conducted in compliance with guidelines the Office of Laboratory Animal Welfare at the National Institutes of Health (United States). C57BL/6 mice aged 6-12 weeks (Jackson Laboratory, USA) were housed in a vivarium and given water and food ad libitum. To obtain vaccinated lymph nodes, male and female mice were subcutaneously vaccinated with 200 μL (50 μL at each flank and shoulder) of Complete Freud’s Adjuvant (InvivoGen) emulsified 1:1 v/v in saline. One week later, lymph nodes were harvested from the mice following humane isoflurane anesthesia and cervical dislocation.

Lymph nodes were sliced according to a previously published protocol.^1^ Briefly, inguinal, axial, and brachial lymph nodes were collected and then embedded in 6 % w/v low melting point agarose (Lonza, Walkersville MD, USA) in 1X PBS. After the agarose had hardened, the section of agarose containing the lymph node was extracted using a 10 mm tissue punch (World Precision Instruments). The agarose block was inverted so that the tissue was at the top of the section and glued to a small mounting stage. The stage was submerged in ice-cold 1X PBS and the lymph nodes sliced into 300-μm thick sections using a Leica VT1000S vibratome (Bannockburn, IL, USA) set to a speed of 90 (0.17 mm/s) and frequency of 3 (30 Hz). Slices were collected and cultured overnight in 500 μL of “complete RPMI”: RPMI (Lonza, 16-167F) supplemented with 10% FBS (VWR, Seradigm USDA approved, 89510-186), 1x L-glutamine (Gibco Life Technologies, 25030-081), 50 U/mL Pen/Strep (Gibco), 50 μM beta-mercaptoethanol (Gibco, 21985-023), 1 mM sodium pyruvate (Hyclone, GE USA), 1x non-essential amino acids (Hyclone, SH30598.01), and 20 mM HEPES (VWR, 97064-362).

### 2.2 Flow Cytometry

Prior to analysis, lymph node slices were removed from the agarose with a paint brush. Slices were crushed individually through a 70-μm nylon mesh filter (Thermo Fisher, USA) by using the rubber tip of a 1- or 3-mL syringe plunger to generate individual cell suspensions. The suspensions were spun down, resuspended in 500 μL of 1x PBS, transferred to a 1.5 mL centrifuge tube, spun down, resuspended in 75 μL of 67 nM Calcein AM (eBioscience) in 1x PBS, and stained for viability for 20 minutes at 37 °C. Stained samples were washed and resuspended in 500 μL of 1x PBS + 2% FBS (flow buffer) before 5 μg/mL of 7-AAD (AAT Bioquest) was added. The samples were run on a Guava 4-color cytometer (6-2L) and analyzed using the Guava^®^ InCyte™ Software. Viable cells were defined as Calcein positive and 7-AAD negative (Figure S1). Viability compensation controls were run on murine splenocytes. 7-AAD controls were run with mixed live and killed cells; cells were killed with 35 % ethanol for 10 minutes at room temperature. Live cells were stained with Calcein-AM for 20 minutes at 37 °C, washed, and mixed with unstained live cells in a 1:1 ratio to act as a single stain compensation control.

### 2.3 ELISA

Slices were stimulated for 18 hours with 10 μg/mL of R848 (Resiquimod, InvivoGen, San Diego, CA) in the culture media, after which supernatant was collected and analyzed by sandwich ELISA for IFNγ secretion. A high-binding plate (Corning Costar 96 well ½ area, #3690; Fisher Scientific) was coated with 1 μg/mL anti-IFNγ XMG1.2 (Biolegend) in PBS overnight at 4°C, then washed. All washing steps were performed in triplicate with 0.05 % Tween-20 in PBS. Wells were blocked for 2 hours with 1 % BSA and 0.05 % Tween-20 (Fisher Scientific) in PBS (block solution). Serial dilutions of recombinant murine IFNγ (Peprotech, Rocky Hill, NJ), were prepared in a 1:1 v/v mixture of block solution and complete media, and supernatant samples were diluted 1:1 v/v with block solution. Samples were added to the plate in duplicate and incubated for 2 hours, then washed. Biotinylated anti-IFNγ R46A2 (0.5 μg/mL, Biolegend) was prepared in blocking solution and added to the plate. Avidin-HRP (1X) (eBioscience) in blocking solution was added to the plate and incubated for 30 minutes, then washed. Plates were developed using TMB substrate (Fisher Scientific), stopped with 1M sulfuric acid (Fisher Scientific), and absorbance values were read at 450 nm on a plate reader (CLARIOstar; BMG LabTech, Cary, NC). To determine concentration of sample solutions, calibration curves were fit in GraphPad Prism 6 with a sigmoidal 4-parameter dose-response curve,

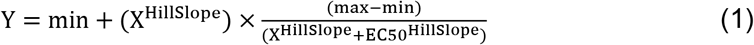

where X is concentration in pg/mL, Y is absorbance (arbitrary units), min and max are the plateaus of the sigmoidal curve on the Y axis, EC50 is the concentration that gives an absorbance value halfway between the minimum and maximum, and HillSlope describes the steepness of the slope. The limit of detection (LOD) was calculated from the concentration corresponding to the average signal of the blank + 6x standard deviation of the blank and was 8.56 pg/mL.

### 2.4 Fluorescent Microsphere Uptake

A microsphere suspension was prepared by diluting 2 μm fluorescent polystyrene microspheres (Fluoresbite^®^ YG Carboxylate Microspheres, Polysciences, 2.5% microsphere suspension) 1:1 (v/v) in flow buffer. After overnight culture, the slices were incubated for 1 hour with 30 μL of the microsphere suspension. After an hour, the slices were washed for one hour in 1x PBS to remove any microspheres that were not uptaken. Slices were imaged on a Zeiss AxioZoom upright macroscope with a PlanNeoFluor Z 1x/0.25 FWD 56mm objective, Axiocam 506 mono camera and HXP 200 C metal halide lamp (Zeiss Microscopy, Germany) using transmitted light (brightfield) and fluorescence in the bead channel (FL Filter Set 38 HE). Image analysis was completed using FIJI software, version 1.53c.^11^ A region of interest was defined by thresholding the outline of the tissue slice in a brightfield image. The mean grey value (MGV) was quantified inside of this region in the corresponding fluorescence image.

### 2.5 DQ-OVA Culture of Lymph Node Slices

Slices were collected as above and randomly assigned to be cultured with either with DQ-ovalbumin (DQ-OVA) protein solution, consisting of 1 μg/mL DQ-OVA (Thermo Fisher, USA) plus 9 μg/mL purified OVA (InvivoGen, USA) in 500 μL complete RPMI, or regular OVA protein solution, containing 10 μg/mL purified OVA in 500 μL complete RPMI. Slices were incubated for 24 hours at 37 °C, 5 % CO_2_, and images were collected at 2, 4, and 24 hours on a Zeiss AxioZoom upright microscope using the Filter set 38 HE. Image analysis was performed FIJI version 1.53c. A region of interest was defined from the brightfield image as in section 2.4, and the MGV within the outline was measured in the corresponding FITC channel.^11^

## 3. Results

The unintentional loss of vaccinated LNs due to their popping out of the agarose selectively biases the slices that can be studied ex vivo. This bias could prevent the ex vivo analysis of the most immunologically active LNs after vaccination, and thus, could limit the understanding of the critical effects that vaccine adjuvants induce that shape adaptive immunity. We identified digitonin as a detergent with the potential to limit LN popping from the agarose after vaccination. To determine the optimal concentration of digitonin, we subcutaneously vaccinated mice with Complete Freud’s Adjuvant (CFA), a highly inflammatory vaccine adjuvant, and harvested the inguinal, brachial, and axial LNs a week later. Harvested LNs were either placed directly into a collection dish containing media, or they were dipped briefly (< 1 sec) into digitonin solution, rinsed with PBS for several seconds, and placed into the collection dish (Figure 1a, Supplementary Video 1). An oily layer of lipids formed atop the digitonin solution after a vaccinated LN was dipped; this layer was not present in the absence of digitonin. The LNs were then embedded in agarose and sliced (Figure 1b) to determine the impact of digitonin on popping (Figure 1c) and slice viability and function.

**Figure 1:**
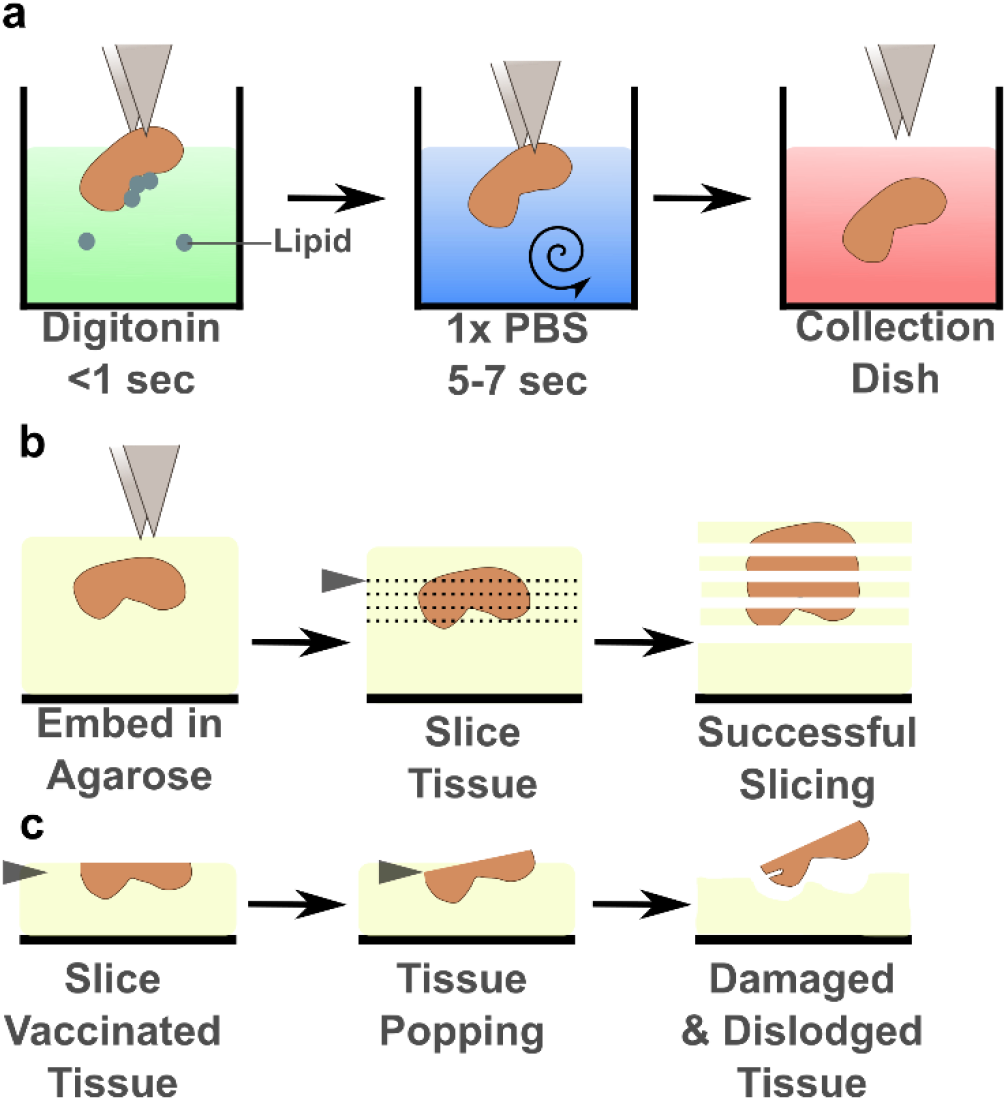
Digitonin wash limited lymph node popping without affecting viability. (a) Schematic of the brief digitonin washing procedure, described in the text. (b) Schematic of successful tissue slicing. The LN is embedded in agarose and sliced on a vibratome (blade, grey triangle) into several 300-μm thick slices that remain adhered to the surrounding agarose. (c) Schematic of LN popping, as the blade approaches the partially sliced organ and dislodges it from the agarose.

The frequency of popping of LNs from vaccinated mice varied drastically between experiments, in some cases being as high as 75% or as low as 14% (Figure 2a). Interestingly, LNs from vaccinated males popped out of the agarose at higher frequencies than female LNs (data not shown). A wash step with 100 μg/mL digitonin significantly decreased the fraction of reactive LNs that popped out of the agarose during slicing (Figure 2a), including in LN from males (Figure S2a). This concentration is below digitonin’s critical micelle concentration of 615 μg/mL. Lower concentrations of digitonin were ineffective in preliminary experiments (Figure S2b). We confirmed that the digitonin wash step did not reduce viability after overnight culture in vaccinated LNs on average (Figure 2c), although there was a small but significant decrease in viability of nodes from vaccinated males (92 vs 89% viability, p = 0.02; Figure S2c). Importantly, digitonin wash also did not decrease viability in slices from naive mice after overnight culture (Figure 2d, Figure S2d). No difference was observed in popping of naïve LNs from the agarose, which occurs infrequently even without digitonin (data not shown).

**Figure 2:**
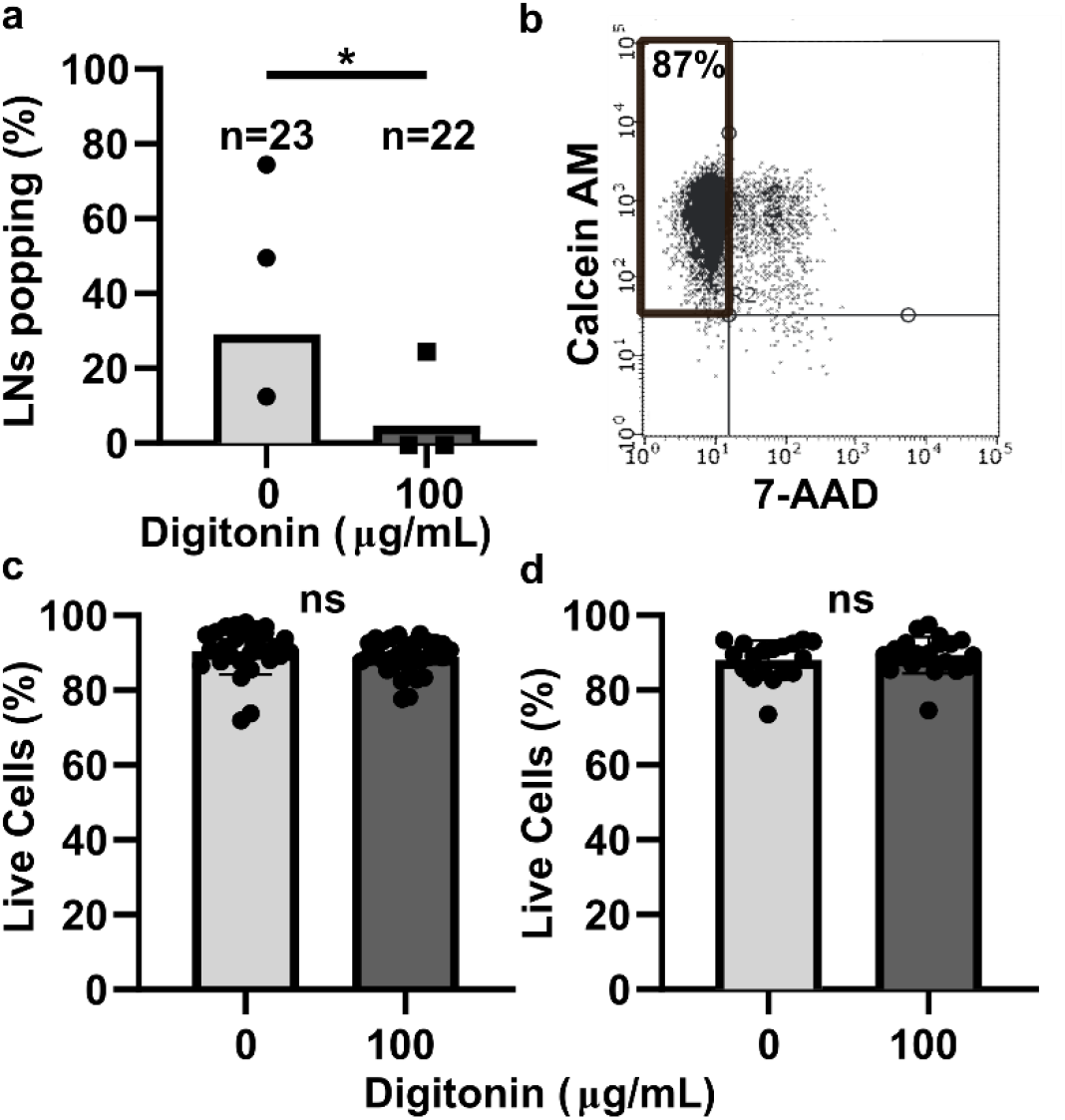
Digitonin wash reduced popping without impacting lymph node slice viability. (a) The percentage of vaccinated LNs that popped during slicing. Each dot represents one experiment. Total n for nodes is shown above each bar. * indicates p=0.0470 by Fisher Exact Test. (b) Representative plot of flow cytometric analysis of viability. Live cells were defined as Calcein AM positive and 7-AAD negative. Data from an unwashed LN from a naïve female mouse are shown. (c,d) Overnight viability of slices from male and female mice that were vaccinated (c) or naïve (d). Each dot represents one slice. Bars show mean and standard deviation. ns indicates p > 0.05 by unpaired t test.

Next, we tested the extent of the digitonin wash’s impact, if any, on specific cell activities in naïve lymph node slices. We reasoned that any digitonin that penetrated the capsule would first enter the peripheral sinus, which are lined with antigen presenting cells (APCs). Therefore, we investigated the extent to which antigen capture and antigen processing by APCs was altered. First, we harvested and sliced LNs from naïve mice and cultured these slices overnight to allow for any delayed toxicity. After overnight culture, 2 μm fluorescent microspheres were added to slices to visualize bead capture. There was no significant difference in the mean fluorescent intensity of the microspheres in slices collected with and without digitonin (Figure 3a), and the distribution of microsphere uptake throughout the tissue was qualitatively similar (Figure 3b), suggesting that penetration and uptake of the beads was not affected at a gross level by the digitonin wash.^12^ Next, we assessed antigen processing by using DQ-ovalbumin (DQ-OVA), which becomes fluorescent as it is endocytosed and proteolytically cleaved. The digitonin wash did not impact antigen internalization and processing; both the rate of uptake and the qualitative distribution were unaltered (Figure 3c, d). Finally, we performed a simple assay of T cell function in the tissues, as this activity is of particular interest in vaccine studies.^1^ Slices were stimulated with R848 overnight and the supernatant analyzed for IFNγ secretion. The digitonin wash step did not significantly alter IFNγ production (Figure 3e), although there was significant variability within each condition. As all slices from the lymph nodes were used, i.e. without pre-selection of a particular depth in the organ, the variability in cytokine secretion is expected and consistent with known heterogeneity in cell number and composition.^1^ In summary, from these data we concluded that the digitonin wash step did not significantly impact APC uptake and antigen processing nor T cell activation, and thus was suitable for use in experiments designed to measure these functions in lymph node slices.

**Figure 3:**
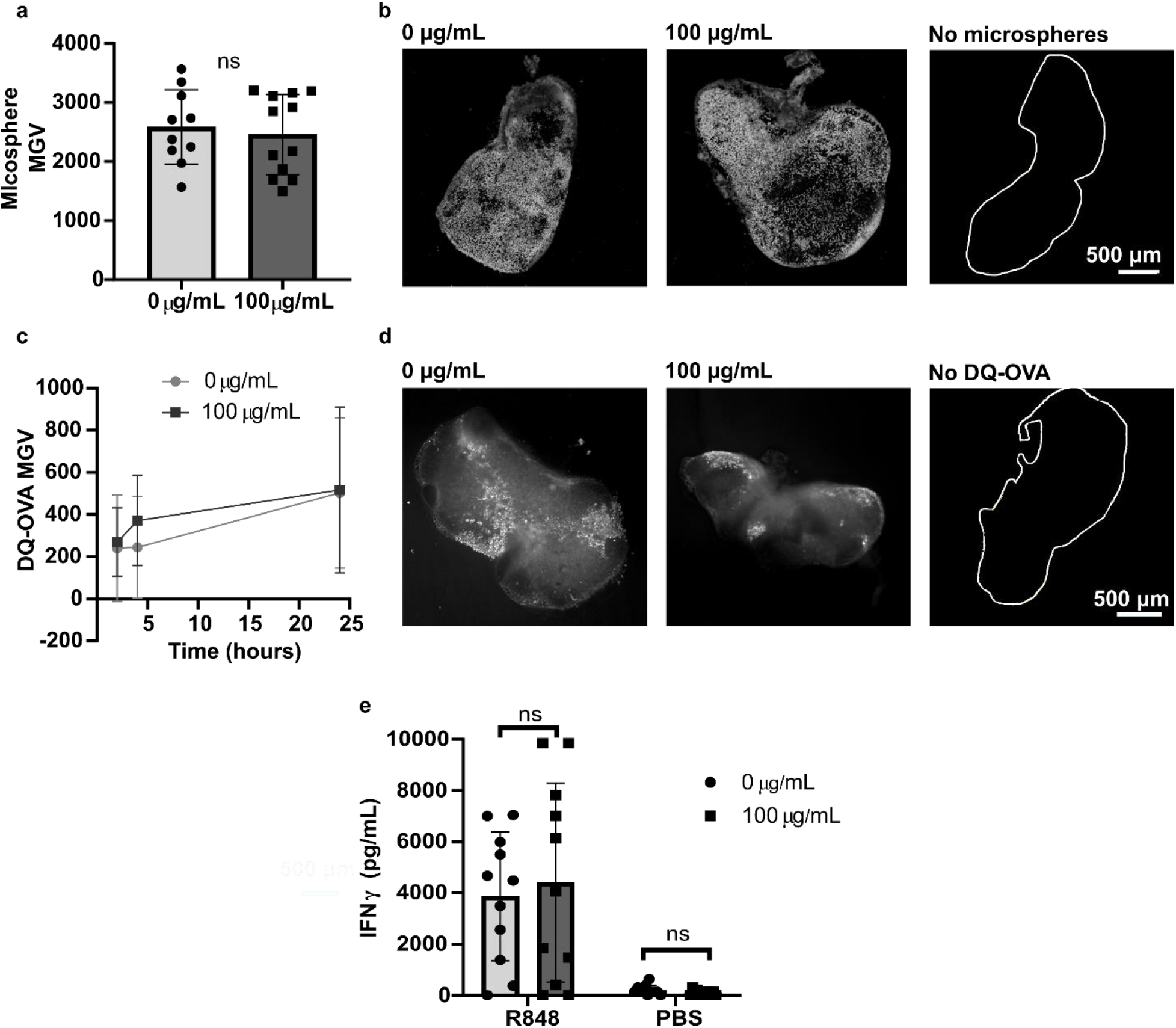
Digitonin wash step does not impact slice function. (a) Mean grey value (MGV) of 2-μm microsphere fluorescence in naïve slices collected with and without a digitonin wash (100 μg/mL), used as a metric of bead uptake. The MGV of each slice was background subtracted by the MGV of a separate set of slices that received no microspheres. ns indicates p > 0.05 by unpaired t test. Each dot represents one slice (n = 10 – 12). Bars show mean with standard deviation. (b) Representative images of microsphere uptake, with and without a digitonin wash. The white trace identifies the outline of a LN slice that received flow buffer instead of microspheres. (c) MGV of DQ-OVA in naïve lymph node slices collected with and without a digitonin wash, used as a metric of antigen uptake, and (d) representative images of DQ-OVA processing at 24 hours. The white trace identifies the outline of a LN slice that received OVA instead of DQ-OVA. Dots show mean and standard deviation. n = 15 for 100 μg/mL; n = 9 for 0 μg/mL. (e) IFNγ concentration in culture supernatant was not significantly different between unwashed and digitonin-washed slices after overnight stimulation with R848. Analyzed by two-way ANOVA. Each dot represents one slice (n = 11). Bars show mean with standard deviation. ns indicates p > 0.05 by unpaired 2-way ANOVA.

## 4. Conclusion

The technical note describes the utility of a digitonin wash step to increase the efficacy of slicing live lymph nodes from vaccinated animals. We found that a brief wash with 100 μg/mL digitonin limited the number of LNs that popped out the agarose during slicing without impacting slice viability or function. Specifically, we determined that the digitonin wash did not impact APC internalization of microscale particles or processing of a protein antigen, nor did it impact a slice’s ability to respond to TLR stimulation. By preventing selective loss of the most reactive lymph nodes, this novel technique will enable robust experiments to study the development of adaptive immune responses in ex vivo LN tissue.

## Acknowledgements

Research reported in this publication was supported by The Hartwell Foundation and by the National Institute of Allergy and Infectious Diseases of the National Institutes of Health under Award Number R01AI131723. The content is solely the responsibility of the authors and does not necessarily represent the official views of the National Institutes of Health. M. C. Belanger was supported in part by the Immunology Training Grant at the University of Virginia (NIH, 5T32AI007496-23). Additionally, the authors would like to thank Megan Catterton, Rebecca Yoo, and Paola Covarrubias for their technical assistance.

## Supporting Information

Supporting information includes a supplemental movie and supplemental figures.

