## Supplemental Information for "Detergent wash improves vaccinated lymph node handling ex vivo"

### Contents

- Supplemental Video Caption
- Supplementary Figures

**Supplementary Video 1:** Movie of washing lymph nodes in digitonin solution. Harvested lymph nodes were quickly dipped into a 100 µg/mL digitonin solution (< 1 sec) before being rinsed in phosphate-buffered saline (3 – 5 sec) and placed into complete RPMI. Movie plays in real time.

**Supplementary Figures:**

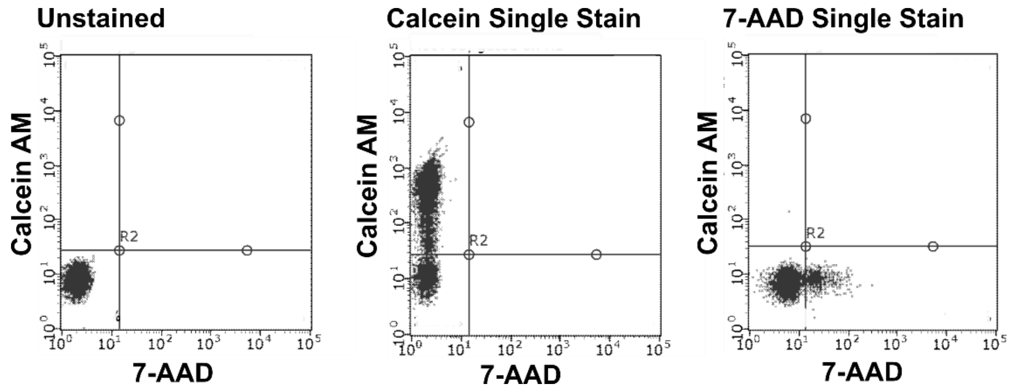


Figure S1: Single stain controls for flow cytometry, as described in Methods.

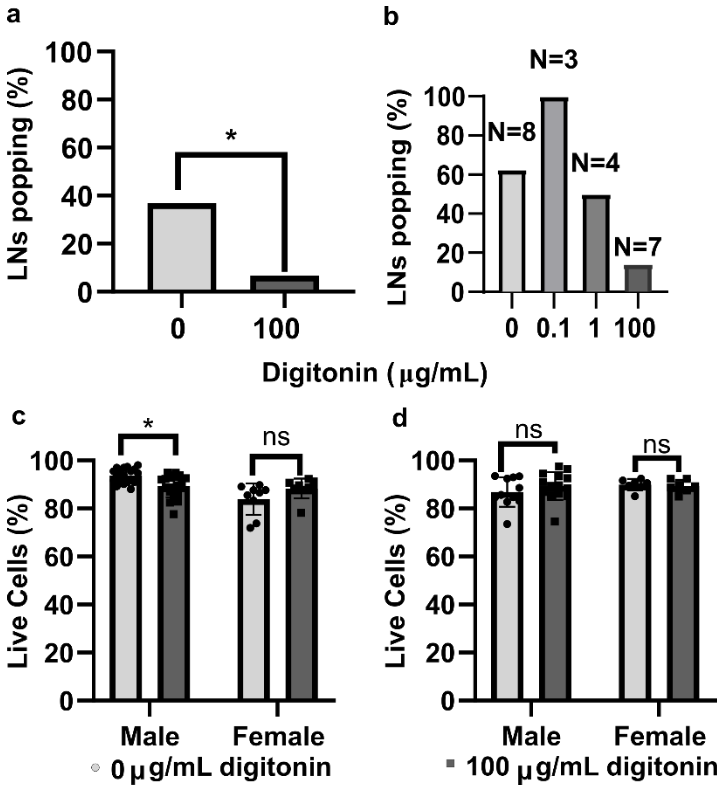


Figure S2: Optimization of digitonin wash and analysis of sex differences in digitonin washing. (a) Popping frequency of LNs from male vaccinated mice with or without digitonin wash. n = 16 for 0 μg/mL, n = 15 for 100 μg/mL. * indicates $p < 0.05$ by unpaired t test. (b) Popping frequencies of LNs from vaccinated animals (male) in preliminary tests with varied concentration of digitonin. (c,d) Overnight viability of lymph node slices with or without a digitonin wash, broken out by sex, for vaccinated mice (c) or naïve mice (d), as assessed by flow cytometry using 7-AAD and Calcein staining. Each dot represents one slice. Bars show mean and standard deviation. * indicates $p < 0.05$ by Two-way ANOVA Multiple Comparisons.
